# A framework for designing miRNA-based distributed cell classifier circuits

**DOI:** 10.1101/2020.05.13.092908

**Authors:** Melania Nowicka, Heike Siebert

## Abstract

**Motivation:** Cell classifiers are synthetic bio-devices performing type-specific *in vivo* classification. The circuits identify a cell state based on its molecular fingerprint. In particular, the classifiers may be designed to recognize cancerous cells and trigger their apoptosis, shaping a novel therapy for cancer patients. Recently, we introduced a new theoretical design of such devices employing distributed classifiers. Here, a group of single-circuit classifiers decides collectively according to a pre-defined threshold function whether a cell is cancerous. The multi-circuit architecture has shown the potential to predict the cell condition with high accuracy. However, lack of far-reaching machinery to design and evaluate distributed cell classifiers, in particular, assessing their robustness to noise and novel information, makes their application limited.

**Results:** In this study, we present a comprehensive framework for designing and evaluating miRNA-based distributed cell classifiers comprising data simulation, pre-processing, and an extensive testing scheme. We develop optimization criteria that allow increasing the accuracy and robustness of classifiers to noise and novel information as shown in simulated and real-world data studies. The evaluation performed on cancer data demonstrates that distributed classifiers outperform single-circuit designs in terms of prediction accuracy. Our classifiers include relevant miRNAs previously described in the literature, as well as more complex regulation patterns included in the data.

**Availability:** The code and data are available at: https://github.com/MelaniaNowicka/RAccoon.

## 1 Introduction

Synthetic biology has shown its potential in the development of therapy-oriented synthetic gene circuits (Xie *et al*., 2011; Ye and Fussenegger, 2014; Kis *et al*., 2015; Slomovic *et al*., 2015). In particular, synthetic switches that convert biological signals into a discrete on/off output may be applied in a diagnostic and therapeutic manner (Xie *et al*., 2011; Slomovic *et al*., 2015). Recently, cell classifier circuits that also follow switch-like behavior are intensively studied (Xie *et al*., 2011; Sayeg *et al*., 2015; Mohammadi *et al*., 2017; Gam *et al*., 2018; Dastor *et al*., 2018). Cell classifiers are synthetic circuits that aim at *in vivo* classification of a cell’s type or current state based on its molecular fingerprint (Xie *et al*., 2011). The circuits are delivered into the cells using, e.g., viral vectors or plasmids. In a cell, the circuit senses type-specific biological signals (inputs) and trigger a controlled production of the desired compound (output) according to a predefined on/off function (Xie *et al*., 2011; Mohammadi *et al*., 2017). The classifiers employ varied molecular signals as inputs, e.g., miRNAs, transcription factors or promoter activities (Dastor *et al*., 2018). Here, we focus on miRNA-based cell classifiers that may selectively target cancer cells by triggering their apoptosis. Inter- and intratumoral heterogeneity poses a major challenge in cancer treatment. Thus, general anticancer therapies are often unsuccessful (Ramón y Cajal *et al*., 2020). Facing cell heterogeneity, patient-tailored methods such as cell classifiers have the potential to be more effective (Xie *et al*., 2011; Dastor *et al*., 2018). Classifier devices may be also applied beyond the described scope, e.g., as cell population purification methods or neuron-type-specific targeting technologies (Sayeg *et al*., 2015; Miki *et al*., 2015). Circuits that employ miRNAs as inputs may be designed based on miRNA differential expression profiles that come from different experiments, e.g., microarray or RNA-seq. However, such experiments do not provide information at the single-cell level. In the future, the data may theoretically be acquired by single-cell miRNA profiling of cancerous and non-cancerous cells collected from a given subject or from *in vivo* cell profiling (Faridani *et al*., 2016; Dastor *et al*., 2018).

Due to the discrete behavior of the circuits, logic modeling is commonly applied to the classifier design (Xie *et al*., 2011; Mohammadi *et al*., 2017; Becker *et al*., 2018). Mohammadi *et al*. (2017) proposed a boolean representation of a classifier, where logic inputs are miRNA expression levels described as low (0) or high (1) and the output is the cell state classified as non-cancerous (0) or cancerous (1). The classifier is represented by a boolean function that captures a relationship between the miRNA regulation and the cell state. According to the output the drug production is activated in the diseased and repressed in the healthy cells. The boolean representation of a classifier is additionally restricted by the assembly constraints, i.e., the classifier should not be too complex and may consist of only a few biological modules that are feasible to construct in the lab (Mohammadi *et al*., 2017). Thus, two main optimization criteria, namely accuracy and simplicity, must be taken into account while designing cell classifiers. Mohammadi *et al*. (2017) proposed an evolutionary algorithm that optimizes a boolean representation of a classifier based on a theoretical biochemical model of the circuit. The parameters of the model are predefined for a given miRNA expression profile. The algorithm optimizes circuit output concentration in healthy and diseased cells. A second computational method was introduced by Becker *et al*. (2018). The authors described a boolean approach employing a logic solver that finds globally optimal classifiers based on discrete data. The algorithm allows optimizing the number of false predictions of a boolean function. When compared employing discrete data, the approach proposed by Becker *et al*. (2018) outperformed the previous method.

The design of cell classifier circuits often requires stable performance in a vacillating environment, posing a major challenge to computational modeling approaches. In particular, the classifiers must demonstrate high robustness to noise and novel information. Also, with the increasing variance in the expression data, the single-circuit classifiers seem to become more complex making their assembly difficult (Mohammadi *et al*., 2017). Recently, we introduced a new theoretical design of cell classifier circuits, employing so-called distributed classifiers (Nowicka and Siebert, 2019). Distributed classifiers were previously described by Didovyk *et al*. (2015) as a population of genetically engineered microbial cells containing simple synthetic biosensors. Such population forms one complex classifier trained to find biological patterns according to a threshold function (Didovyk *et al*., 2015; Kanakov *et al*., 2015). In terms of cancer cell classification, the distributed classifier (DC) is a set of small single-circuit classifiers that decides collectively based on a threshold function whether the drug production should be activated or repressed in a cell (Nowicka and Siebert, 2019). The threshold-based decision scheme allows for a reduction of false predictions caused by errors of individuals. Thus, the DC design may allow increasing the performance of classifiers when facing the heterogeneity of the cell environment. Mohammadi *et al*. (2017) mentioned that such independent circuits operating simultaneously may be also more robust in terms of tumor mutational escape mechanism (Mohammadi *et al*., 2017). In the previous work, we proposed a genetic algorithm (GA) that allows designing DCs described in detail in Nowicka and Siebert (2019). The real-world case study performed on a breast cancer data set shows that DCs may achieve higher accuracy than single-circuit classifiers. Although the study demonstrated the potential of distributed cell classifiers, neither the algorithm nor the testing scheme went beyond a proof-of-concept.

Our goal in this work is to develop the mentioned algorithm into a comprehensive and accessible framework for the design of distributed cell classifiers by addressing core needs in the application. We propose a design-independent workflow for data simulation and processing as well as evaluation of cell classifiers employing synthetic and real-world cancer data. Previously, such machinery was not available (Mohammadi *et al*., 2017; Becker *et al*., 2018; Nowicka and Siebert, 2019). The design of cell classifiers begins with data processing that facilitates capturing truly differentially expressed miRNAs. First, the raw data should be processed and normalized according to the requirements of a given experimental platform (Soneson and Delorenzi, 2013). Then, to provide clear-cut information about the miRNA regulation across different samples the miRNA expression profiles can be discretized. Such data transformation was shown to improve classification accuracy when applied to biomedical data, in particular, cancer gene expression data (Lustgarten *et al*., 2008; Wang *et al*., 2014; Gallo *et al*., 2016). However, it is important to choose the discretization method tailored to the needs of data and given classification problem. In this study, we employ a discretization approach proposed by Wang *et al*. (2014) that has been shown to outperform other widely-used approaches in terms of cancer classification. The method allows computing the overall capability of a gene to distinguish between the classes. We employ this information as a measure of miRNA robustness and propose a multi-objective function that allows measuring the classification performance of DCs and the quality of used miRNA inputs.

Similarly to the optimization process, the evaluation of classifiers should be performed with the emphasis on their robustness to different types of variance in the profiles and novel information not present in the training data. Exact measurements that allow capturing all the different patterns of miRNA expression are not available due to cell abundance and heterogeneity. Thus, the classifiers must have built-in robustness to different scenarios. Previously mentioned methods apply limited testing schemes to assess the classifier’s performance, e.g., do not employ holdout testing data (Mohammadi *et al*., 2017; Becker *et al*., 2018; Nowicka and Siebert, 2019). Extending the evaluation scheme beyond cross-validation provides a reliable estimation of the generalization error.

In this article, we present a comprehensive framework for designing and evaluating distributed classifiers including data simulation, preprocessing and an extensive testing scheme. To increase the built-in robustness of classifiers we introduce a tunable multi-objective function taking the quality of employed miRNAs into account. Further, we develop a testing scheme that allows assessing the generalization ability of classifiers while facing noise and novel information. We assess DC’s performance by employing a fully controllable setting by generating classifier-independent synthetic data sets introducing different types of variance. Then, we evaluate the performance of the new multi-objective function in comparison to the one previously applied in Nowicka and Siebert (2019). Finally, we use cancer data sets coming from studies employing different experimental platforms to assess the robustness of classifiers on real-world cancer miRNA expression profiles and compare the performance of single-circuit and distributed classifiers. Last, we present an exemplary distributed classifier and analyze its architecture.

## 2 Preliminaries

### 2.1 miRNA expression data

MicroRNAs are differentially expressed between different cell types, as well as different cell conditions, in particular, between cancerous and non-cancerous cells. Thus, dysregulated miRNAs have been considered diagnostic and prognostic biomarkers in cancer (Lan *et al*., 2015). This allows capturing the differences between healthy and cancerous tissues and classifying them based on a type-specific miRNA profile. In this work, we employ simulated RNA-seq and real-world cancer data coming from different platforms. All the data sets are initially transformed into a format presented in Table 1. The first column includes unique sample IDs and the second its annotation, where 0 stands for non-cancerous and 1 for cancerous cells. The annotation is followed by a miRNA expression profile consisting of unique miRNA IDs and their expression levels in each sample. Such data sets may include different types of information, e.g., RNA-seq counts or microarray measurements.

**Table 1.** Exemplary miRNA profile.

| ID | annotation | miR-a | miR-b | miR-c |
| --- | --- | --- | --- | --- |
| 1 | 0 | 126 | 629 | 21 |
| 2 | 0 | 35 | 4664 | 979 |
| 3 | 1 | 744 | 3728 | 32 |
| 4 | 1 | 888 | 7068 | 11 |

### 2.2 Discretization of data

To discretize the miRNA expression profiles we employ the method proposed by Wang *et al*. (2014). The authors demonstrated that their approach outperforms other commonly used supervised discretization methods when applied to the classification of cancer gene expression data (Wang *et al*., 2014; Gallo *et al*., 2016). The described method allows discretizing expression of each miRNA in the data into one, two or three states by finding zero, one or two cut-points, respectively. This allows describing the expression pattern across all the samples. The miRNA may be (i) non-regulated, (ii) differentially regulated between the classes or (iii) follow a more complicated regulation pattern, e.g., be both up- and down-regulated in one class and non-regulated in the other class. In the latter case, a miRNA is discretized into 3 states: −1, 0 and 1 being not compatible with the boolean representation of a classifier. Thus, in terms of cell classifier application, miRNAs that follow the second pattern (ii) are candidates for the inputs.

To find the thresholds, the expression range of each miRNA is divided into *m* (*m* ⩾ 50) left-side-half-open intervals, where *V_i_* = (−∞, *l_i_* is the i-th half-open interval and *l_i_* = 1, 2,…, *m* are the upper-boundaries of the intervals. For each of the intervals the class distribution diversity CDD(vi) is calculated according to Eq.1:

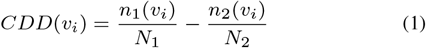

where *n*_1_ (*v_i_*) and *n*_2_ (*v_i_*) represent the numbers of samples belonging to the first or the second class in the interval *v_i_*. *N*_1_ and *N*_2_ are the total numbers of samples in each class. Then, among all the intervals, two with the maximum and minimum CDDs (CDD_max_ and CDD_min_) are found and the global class distribution diversity of a miRNA (Δ) may be

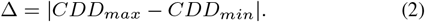

Here, Δ describes the overall capability of a feature to distinguish between two classes. We employ this measure to assess the robustness of miRNAs as classifier inputs (described further in section 3.4). The authors employ Δ, *CDD_max_*, *CDD_min_* and two other tunable parameters *α* and *λ* to find the discretization cut-points based on 3 criteria described in details in Wang *et al*. (2014). In this study, we refer to the discretization parameters as *m_bin_, α_bin_* and *λ_bin_* By increasing *α_bin_* value one may raise the lower bound for Δ to define a miRNA as differentially regulated. Using those criteria we transform the continuous data described in the previous section into discrete profiles where the miRNAs are either differentially regulated between the classes (ii) or non-relevant (i and iii). This allows to transform the data set presented in section 2.1 into binarized data sets *D* = (*S, A*) where *S* ⊆ {0, 1}^*m*^ is the set of sample vectors, 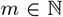 is the number of miRNAs and *A* : *S* → {0, 1} is the sample annotation. Non-relevant features may be removed from further analysis, which allows increasing the computation efficiency as the data size may be significantly reduced.

### 2.3 Single-circuit classifier

A single-circuit classifier can be represented by a boolean function *f* : *S* → {0,1}, where logic inputs are miRNA expression levels defined as low (0) or high (1) and the output is the cell condition classified as non-cancerous (0) or cancerous (1). Due to the lab assembly limitations, the function must be given in the *Conjunctive Normal Form* (Mohammadi *et al*., 2017). This representation corresponds to the combination of the available biological building blocks. A CNF function is a conjunction of clauses where each clause is a disjunction of negated or non-negated literals (inputs). In terms of classifier design, the literals correspond to the miRNA and the clauses to the logic gates. The assembly restrictions pose further constraints on the function. Ultimately, the function may consist of up to 10 miRNA inputs and 6 different gates in total. Two gate modules were proposed by Mohammadi *et al*. (2017): an OR gate with up to 3 inputs and a NOT gate with 1 input. In terms of gates, the circuit may consist of up to 2 OR gates and up to 4 NOT gates. The following function: (*miR-a OR miR-b*) *AND*(*NOT miR-c*) is an example of a 3-input classifier consisting of 2 gates. The function outputs 1 (activation) if either *miR-a* or *miR-b* is up-regulated (1) and *miR-c* is down-regulated (0).

### 2.4 Distributed Classifier

A Distributed Classifier (DC) is a finite set *DC* = *{f_1_,…, f_c_*}, where *c* is the DC’s size and each *f_i_* : *S* → {0, 1} is a unique single-circuit classifier (rule). To simplify the design of SC classifiers the number of unique miRNAs that may be inputs is restricted to 2. Also, to reduce the overall complexity of DC’s architecture and to compare the performance of both, single-circuit and distributed classifier designs the number of rules is restricted to 5 (Nowicka and Siebert, 2019). This limits the maximal number of inputs in a DC to 10 as proposed for SC classifiers by Mohammadi *et al*. (2017). An example of a DC is presented below.

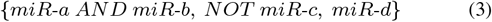

DCs classify cells as non-cancerous (0) or cancerous (1) according to a threshold function *O_DC_* : *S* → {0,1}:

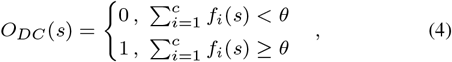

where *s* ∈ *S* is a sample, *θ* = [*α · c*⌉ is a decision threshold (rounded half up), 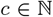 is the classifier size and *α* is the ratio that allows calculating the decision threshold based on the DC’s size. The *α* value corresponds to the minimal fraction of rules in the DC that must output 1 (cancerous) for the DC to classify a cell as cancerous. A cell is correctly classified by a given DC if the classifier’s output agrees with its annotation. Considering the example presented in Eq. 3 (i) for low *α* (*θ* =1) only one rule must output 1 for the DC to output 1, (ii) for intermediate *α* (*θ* = 2) two of the rules, (iii) for high *α* (*θ* = 3) all of the rules. In the first case, the function becomes a disjunction of the rules. In the second, the DC decides according to a verdict of the majority. In the last case, the function becomes a conjunction of rules being similar to a large single-circuit classifier. Thus, the classifier’s output may substantially differ for varying values of *α*. This may influence the sensitivity and specificity of the classifiers as shown in Nowicka and Siebert (2019).

## 3 Methods

### 3.1 Data simulation and pre-processing

#### 3.1.1 Data simulation

Data simulation provides a controllable environment to evaluate the classification performance of a new method facing pre-defined experimental conditions. Here, we set up an evaluation strategy to test distributed classifiers against the noise of different intensity and type. Six synthetic RNA-seq data sets were generated using the compcodeR package developed by Soneson (2014). CompcodeR is a benchmarking Bioconductor package that allows simulating differential gene expression RNA-seq experiments using various parameters. This allows us to fully control the setting of each experiment in regard to the number of samples in each class and the fraction of truly differentially regulated genes, as well as the level of noise in the data (Soneson and Delorenzi, 2013). As a default, compcodeR uses Negative Binomial distribution to simulate RNA-seq counts. The parameters of the distribution, the mean and the dispersion are estimated from real-world data sets (Soneson and Delorenzi, 2013). We generated the data using the following parameters: (i) 200 samples in total, 100 samples per class, (ii) 1000 genes, of which 10% are differentially expressed, (iii) 50% of the differentially expressed genes are up-regulated. The number of samples and features approximately corresponds to the maximal size of cancer data sets employed in this study (described in Section 3.2.1). The fraction of differentially expressed and up-regulated genes come from the setting proposed by Soneson and Delorenzi (2013).

To test the robustness of the DCs we introduced different types of variations in the data. First, we tested the classifiers against the occurrence of random outlier values in the data. In this case, each observed count is multiplied or divided by a random number between 5 and 10 with equal probability of the value being increased or reduced. We introduced the outliers with probabilities varying between 0.0 and 0.3 with step 0.1 labelled as SDR0, SDR10, SDR20 and SDR30, where the number determines the probability of random outliers allowed in the simulation. All the other parameters were set to default.

We also tested two other scenarios by altering the distribution parameters. First, we generated a data set with different dispersions between the two conditions using the Negative Binomial distribution (SDDISP). This allows to evaluate the DC’s performance against different spread of expression values between the classes. Then, we generated a data set where 50% of the genes were simulated according to a Negative Binomial distribution and 50% according to Poisson distribution (SDP50) to assess the DC’s robustness against features following different distributions across the samples. After simulation, each of the data sets was split into 80% training and 20% testing fractions.

#### 3.1.2 Normalization

The synthetic data sets were normalized with the TMM (trimmed mean of M-values) normalization method available in the edgeR package (Robinson *et al*., 2009). The normalization strategy is based on an assumption that most of the genes in the data are not differentially expressed (Dillies *et al*., 2013). The normalization factors are calculated using a reference sample chosen from all the samples in the data set (Robinson *et al*., 2009). TMM normalization was shown to maintain low false-positive rate in terms of detection of differentially regulated genes while facing count variations in the data (Dillies *et al*., 2013). The training and testing data sets were normalized separately to avoid information leakage. The reference sample was chosen based on the training data set and then used to normalize both, the training and testing data sets.

#### 3.1.3 Discretization

The data sets were binarized using the discretization method by Wang *et al*. (2014) described in section 2.2. The training and testing data sets were discretized separately. First, the discretization thresholds for each feature in the data set were calculated based on the training samples with the default parameters proposed by Wang *et al*. (2014): *m_bin_*=50, *α_bin_*=0.5, *λ_bin_*=θo1o Then, the thresholds were applied to the discretization of the testing data set. All the features that were non-regulated or followed a 2-cut-point discretization pattern were filtered out from the training data. For each feature in the training data the global class distribution diversity Δ was calculated and stored.

### 3.2 Case studies data

#### 3.2.1 Cancer data sets

We employed the dbDEMC 2.0 database (Yang *et al*., 2017) to acquire cancer miRNA expression profiles that follow cancer vs. control design. We searched for balanced data sets, i.e., data sets containing an equal number of samples representing both conditions to avoid bias towards one of the classes (Yang *et al*., 2006). Finally, 3 real-world cancer data sets were downloaded from the NCBI’s Gene Expression Omnibus (Edgar, 2002). The details regarding each of the data sets employed in this study are presented in Table 2.

**Table 2.** Cancer data sets description.

| GEO Acc. No. | cancer type | samples | per condition | miRNA | filtered |
| --- | --- | --- | --- | --- | --- |
| 22058 | HCC | 192 | 96 | 220 | 65 |
| 10694 | HCC | 156 | 78 | 110 | 10 |
| 36681FF | NSCLC | 112 | 56 | 858 | 56 |
| 36681FFPE | NSCLC | 94 | 47 | 858 | 19 |

Two hepatocellular carcinoma (HCC) data sets (data accessible at NCBI GEO database (Edgar, 2002): GSE10694 and GSE22058, platform GPL10457), as well as the non-small-cell lung carcinoma (NSCLC) data set (data accessible at NCBI GEO database (Edgar, 2002): GSE36681), come from paired design experiments, i.e., from each subject two samples were collected, one from the cancerous and one from the adjacent non-cancerous tissue (Li *et al*., 2008; Burchard *et al*., 2010; Jang *et al*., 2012). GSE10694 includes 156 paired samples and 10 normal liver samples. The NSCLC data set consists of two subsets separated by the sample preparation procedure (FF - fresh frozen, FFPE - formalin-fixed, paraffin-embedded).

#### 3.2.2 Pre-processing

All the data sets were pre-processed by the authors of the experiments using adequate methods of normalization for applied miRNA profiling platforms. Further, we removed all non-human miRNAs from each of the data sets. Additionally, 10 samples coming from normal liver tissues collected from other subjects (not paired) were removed from GSE10694 data set to maintain an equal number of samples in each class. Both HCC data sets (GSE22058 and GSE10694) were divided into training (80%) and testing (20%) fractions by subject, i.e., both samples, cancerous and non-cancerous from the same subject were randomly added to either training or testing data set. The NSCLC data set (GSE36681) was split into two subsets: GSE36681-FF and GSE36681-FFPE according to the applied sample preparation method. Further, both data sets were randomly divided into training (80%) and testing (20%) fractions as the information about sample pairing was not available. The data sets were discretized analogously to the procedure described for synthetic data. The numbers of miRNAs included in the data sets before and after the selection of relevant miRNAs can be found in Table 2.

### 3.3 Genetic Algorithm

As a starting point, we used the GA architecture applied to the DC problem as described in Nowicka and Siebert (2019). Building on that, we investigated the key aspect shaping the performance of the method, namely the objective function. We introduced multi-objective optimization of classifiers tailored to the demands of the application taking not only accuracy but also robustness into account. The new objective function is described in the following section. Furthermore, to increase the performance of the algorithm we refined the termination criterion. Previously, the number of algorithm’s iterations was restricted by a pre-defined number of iterations. Currently, the termination criterion is specified by the maximal number of last iterations in which the classifier’s performance was not improved.

### 3.4 Objective Function

We apply balanced accuracy (BACC) to evaluate the accuracy of distributed classifiers calculated according to Eq.5:

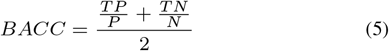

where *TP* and *TN* are the numbers of true positives and true negatives. *P* and *N* are the numbers of the samples in the data representing the positive and negative class. Balanced Accuracy normalizes the correct positive and negative predictions by the number of positive and negative samples in the data set. This allows evaluating the binary classification performance of DCs taking data imbalance into account.

However, miRNAs discretized into two states, i.e., differentially regulated, usually differ in terms of robustness. This can be measured using the global class distribution diversity Δ described in section 2.2. The miRNAs that demonstrate high Δ display more discriminative regulation pattern between the classes. Thus, such miRNAs may be more robust against noise and novel information that is not contained in the training data. Here, we introduce the second measure of classifier performance *DC*_Δ_ calculated according to Eq.6:

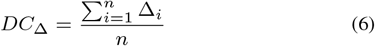

where *i* =1, …,*n* is the ith miRNA in the DC. *DC*_Δ_ describes the average ability of miRNAs in the classifier to distinguish between the two classes. Here, we propose a weighted multi-objective function that allows calculating a combined *DC_score_*:

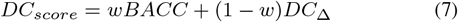

where weight *w* ∈ [0.0,1.0] determines the importance of each objective, i.e., high w emphasizes the importance of BACC and low w the importance of *DC*_Δ_ in the optimization process. The weight allows adapting the significance of both objectives to the data and increases the performance of classifiers in terms of their robustness to noise and novel information. Note, w = 0.0 optimizes only for *DC*_Δ_-Thus, the information about the accuracy of a classifier is not included in the optimization process. In the case of a data set in which miRNAs significantly differ in terms of robustness (Δ), *DC*_Δ_ allows capturing better candidates for classifier inputs. Here, one may expect a higher contribution of the *DC*_Δ_ in the optimization process. Note, w = 1.0 corresponds to the prior objective function proposed by Nowicka and Siebert (2019). In this study, we apply *DC_score_* to train classifiers and BACC to evaluate their performance.

Another important objective in cell classifier design is the simplicity of the circuit. As described in Nowicka and Siebert (2019), the proposed algorithm records all the best solutions found during the GA iterations. After the termination, a classifier with the lowest number of inputs is returned from a pool of equally well-performing solutions.

### 3.5 Test design

We performed tests on simulated and cancer data to measure the performance of DCs in a controlled and real-world setting. As described before, each data set was first divided into training (80%) and testing (20%) fractions and processed according to the description in Section 3. Further, the overall test design was identical for each of the data sets.

The training data set was employed to tune the standard GA parameters: termination criterion (tc), population size (*ps*), tournament size (*ts*), crossover (cp) and mutation probability (*mp*), as well as the objective function weight (*w*). To optimize the parameters of the algorithm (*w*, *tc*, *ps*, *ts*, *cp* and *mp*) we applied a random search approach. Random search allows to obtain performance similar to the grid search approach, while significantly decreasing the computational cost of the tuning (Bergstra and Bengio, 2012). We randomly chose 100 sets of parameters in the following ranges: *w*: 0.1-1.0, step 0.1, *tc*: 10-30, step 5, *ps*: 50-300, step 50, *ts*: 0.1-0.5, step 0.1, *cp*: 0.1-1.0, step 0.1, *mp*: 0.1-1.0, step 0.1.

We performed 10-fold cross-validation (CV) for each generated parameter set. First, we divided the non-discretized training data into 10 pairs of training (90%) and validation data sets (10%), and discretized them analogously to the procedure described in Section 3.1.3. The classifiers were trained on the training data using DC_score_ as the objective function and evaluated on the validation data using BACC. Each single GA run was repeated 10 times and the average score was recorded. We ran the parameter tuning separately for 5 different decision thresholds *α* = {0.25, 0.45, 0.50, 0.75, 1.00} covering all possible *θs* for different sizes of DCs. This allows choosing the best parameter set for each *α*. Then, we selected the parameter set and the best performing *α* based on the average validation BACC resulting from the 10-fold CV. We applied the chosen parameters to train classifiers on the discretized training data set and evaluated the models on the testing data set. We repeated the training 10 times and recorded average scores.

### 3.6 Implementation

The data simulation and pre-processing scripts are implemented in R (except for binarization). The binarization procedure is implemented in Python based on the algorithm proposed by Wang *et al*. (2014). The genetic algorithm and a fully automated testing procedure described in the previous section is implemented in Python. One may choose between running a complex analysis according to the described testing scheme and using the algorithm separately to train the classifiers for a given parameter set. The user may define all the parameters employed in this study or use a pre-defined configuration.

## 4 Results

### 4.1 Simulated data studies

#### 4.1.1 Parameter tuning

The best performing parameter sets found in the tuning process for each of the simulated data sets are presented in Table 3. As described in Section 2.4, the *α* value corresponds to the minimal number of single-circuit classifiers that must agree to classify a cell as cancerous in regard to the DC’s size. We tuned the parameters separately for five different *α* values (0.25, 0.45, 0.50, 0.75 and 1.00). The best performing *α*s differ between the data sets simulated with different parameters, i.e., the decision threshold seems to be sensitive to the changing distributions and intensity of noise in the data. For most of the data sets, intermediate values of *α* (around 0.50) achieved a higher accuracy. Note that *α* is related to the classifier’s size, e.g., for a classifier consisting of only one rule *α* ⩾ 0.50 results in identical threshold θ =1 and for a classifier consisting of five rules the threshold θ differs for all tested *α*s. Thus, *α* values should be interpreted in the context of the classifier’s size.

**Table 3.** Results of parameter tuning for the simulated data sets: *α* - threshold ratio, *w* - objective function weight, tc - termination criterion, ps - population size, cp - crossover probability, mp - mutation probability, ts - fraction of population chosen for tournament selection.

| dataset | $\alpha$ | $w$ | $tc$ | $ps$ | $cp$ | $mp$ | $ts$ |
| --- | --- | --- | --- | --- | --- | --- | --- |
| SDR0 | 0.75 | 0.8 | 20 | 300 | 0.4 | 1.0 | 0.1 |
| SDR10 | 0.45 | 0.9 | 30 | 250 | 1.0 | 0.6 | 0.4 |
| SDR20 | 0.50 | 0.8 | 20 | 300 | 0.4 | 1.0 | 0.1 |
| SDR30 | 0.50 | 0.8 | 20 | 300 | 0.4 | 1.0 | 0.1 |
| SDDISP | 0.50 | 0.9 | 10 | 100 | 0.9 | 0.7 | 0.3 |
| SDP50 | 1.00 | 0.3 | 30 | 50 | 0.5 | 0.2 | 0.4 |

For almost all data sets the weight value varies between 0.8 and 0.9, i.e., the relevance of BACC in the optimization process is higher than *DC*_Δ_. For the last simulated data set the weight drops to 0.3. Here, the *DC*_Δ_ score seems to be a more valuable objective. This may be caused by the different distributions underlying the data influencing the variance in the robustness. The weight did not reach its maximal value (*w*=1.0) for any of the simulated data sets. The results of the parameter tuning suggest that *DC*_Δ_ is significant in the optimization of classifiers and allows to increase their robustness. We analyze the weight performance further in detail using the real-world data.

The tuned GA parameter values vary between simulated data except for three SDR data sets. For SDR0, SDR20, and SDR30 all the parameters are identical. The data sets were simulated according to the same distribution with an increasing probability of random outliers. Thus, the parameters seem to be adjusted to the underlying structure of data. The most outlying parameter set is the one tuned for SDP50 data set. In particular, the chosen population size is significantly lower than for other data sets. Here, 50% of features in the data set follow the Poisson distribution instead of the default Negative Binomial distribution. The parameters seem to be particularly sensitive to such substantial divergence between the data sets. The termination criterion varies between its minimal (10) and maximal (30) value. The range of *tc* values may be extended to ensure the best performance of the algorithm. The average time of single training run for presented parameters tuned for simulated data is 24.52 seconds.

#### 4.1.2 Evaluation

Here, we present the results of DC’s performance recorded for the simulated data sets. We also interpret the meaning of *α* in terms of the size of trained classifiers. Table 4 contains the scores for best performing decision thresholds as well as the average number of rules and inputs in the trained classifiers.

**Table 4.** Performance of DCs for simulated data sets: *α* - threshold ratio, *w* - objective function weight, BACC_cv_ - average CV BACC, BACC_tr_ - average training BACC, BACC_te_ - average testing BACC, *σ*_te_ - standard deviation for BACC_te_, 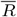 - average number of rules, 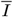 - average number of inputs.

| dataset | $\alpha$ | $w$ | $BACC_{cv}$ | $BACC_{tr}$ | $BACC_{te}$ | $\sigma_{te}$ | $\bar{R}$ | $\bar{I}$ |
| --- | --- | --- | --- | --- | --- | --- | --- | --- |
| SDR0 | 0.75 | 0.80 | 98.92 | 100.00 | 100.00 | 0.00 | 1.0 | 1.0 |
| SDR10 | 0.45 | 0.90 | 96.52 | 100.00 | 98.75 | 1.25 | 3.4 | 4.7 |
| SDR20 | 0.50 | 0.80 | 94.52 | 100.00 | 99.00 | 2.00 | 4.9 | 6.0 |
| SDR30 | 0.50 | 0.80 | 91.45 | 100.00 | 93.50 | 3.74 | 5.0 | 6.4 |
| SDDISP | 0.50 | 0.90 | 98.15 | 99.94 | 100.00 | 0.00 | 1.5 | 2.5 |
| SDP50 | 1.00 | 0.30 | 99.81 | 100.00 | 99.75 | 0.75 | 1.0 | 1.0 |

The results of 10-fold cross-validation show that the optimized DCs generalize well on the validation data (BACC_cv_ above 91% for all data sets). The CV-based tuning provided parameters that allow optimization of classifiers that achieve 100% accuracy for nearly all training data sets, i.e., *BACC_tr_* slightly drops below 100% only for SDDISP. For SDR0 data set (no outliers) the algorithm finds solutions that reach 100% accuracy for both, the training and testing data set. With the increasing probability of random outliers (SDR0-SDR30), the accuracy on validation and testing data declines together with the performance stability (*σ*_te_). However, the BACC_te_ for all data sets is higher than 93%. For the SDDISP data set (different dispersion between classes) as well as for SDP50 (different distributions between samples), the DCs predict the condition with accuracy ⩾ 99.75 for the holdout data. The results demonstrate that distributed classifiers cope well with the different intensity and source of noise in the data.

For two data sets (SDR0, SDP50), the classifiers consist of only one rule on average (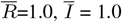, *α* ⩾ 0.75) being in fact 1-input single-circuit classifiers deciding according the verdict of only one rule. However, the average size of classifiers in terms of rules and inputs grows with the increasing intensity of noise in the data (SDR0-30). Simultaneously, the *α* value drops to 0.45-0.50. In regard to the average classifier’s size (3.4 ⩽ 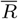 ⩽ 5.0) approximately half of the SC classifiers in a DC must output 1 to classify a sample as cancerous. This shows that distributed classifiers display higher robustness than simple single-circuit classifiers. The decision threshold, as well as the classifier’s size, seem to be adapted to the intensity and source of noise in the data. Thus, the *α* should be tuned for a given data set to increase the accuracy and robustness of classifiers.

### 4.2 Cancer data studies

#### 4.2.1 Parameter tuning

Table 5 includes the parameter sets tuned for real-world cancer data. For most of the data sets the best performing *α* values are equal (*α* = 0.45 for 10694, 36681-FF and -FFPE), i.e., approximately half of the single-circuit classifiers must agree on the classification of cancerous cells. This suggests that distributed classifiers with intermediate-*α* thresholds are suitable for processing real-world cancer data. The *α* is lower only for 22058 data set.

**Table 5.** Results of parameter tuning for real-world data: *α* - threshold ratio, w - weight, *tc* - termination criterion, *ps* - population size, *cp* - crossover probability, *mp* - mutation probability, ts - fraction of population in tournament selection.

| dataset | $\alpha$ | $w$ | $tc$ | $ps$ | $cp$ | $mp$ | $ts$ |
| --- | --- | --- | --- | --- | --- | --- | --- |
| 22058 | 0.25 | 0.20 | 30 | 300 | 0.3 | 0.3 | 0.2 |
| 10694 | 0.45 | 0.80 | 20 | 300 | 0.4 | 1.0 | 0.1 |
| 36681FF | 0.45 | 0.40 | 25 | 150 | 0.1 | 0.7 | 0.5 |
| 36681FFPE | 0.45 | 0.40 | 25 | 150 | 0.1 | 0.7 | 0.5 |

The chosen weight is also low for most of the data sets, in particular, for 22058 data set (*w*=0.2). In those cases, the contribution of the robustness score *DC*_Δ_ is more important to the optimization process. This may suggest that many miRNAs in the data sets, although distretized into two states, differ in their robustness. Here, the *DC*_Δ_ allows to choose better candidates for classifier inputs. All the parameters tuned for the lung cancer subsets (36681-FF and 36681-FFPE) are identical. The data sets come from the same experimental platform and cancer type. Thus, the underlying structure of the data may be very similar. The parameters seem to be again data dependent. Thus, tuning performed for each data set separately is recommended. Average time of single training run for presented parameters tuned for cancer data sets is 18.53 seconds.

#### 4.2.2 Weight evaluation

To evaluate the performance of the proposed multi-objective function we set *α* to the values derived from the parameter tuning described in the previous section: 0.25 for 22058 data set and to 0.45 for the rest of the data set. We re-run tests for 10 weights separately by fixing their values between 0.1 and 1.0 with 0.1 step. We re-tuned the GA parameters for each weight as described in Section 3.5. Table 6 contains the results obtained for the previously tuned weights (Section 4.2.1) and for *w*=1.0 which corresponds to the prior objective function (BACC) applied in Nowicka and Siebert (2019).

**Table 6.** Evaluation of multi-objective function performance: *w* - weight, BACC_cv_ - average CV BACC, BACC_tr_ - average training BACC, BACC_te_ - average testing BACC, *σ*_te_ - standard deviation for BACC_te_.

| dataset | $w$ | $BACC_{cv}$ | $BACC_{tr}$ | $BACC_{te}$ | $\sigma_{te}$ |
| --- | --- | --- | --- | --- | --- |
| 22058 | 0.2 | **98.31** | 97.37 | **97.25** | 0.75 |
|  | 1.0 | 96.31 | **100.00** | 90.00 | 5.00 |
| 10694 | 0.8 | **87.11** | **94.35** | **85.31** | 1.43 |
|  | 1.0 | 86.66 | 94.27 | 84.38 | 3.42 |
| 36681FF | 0.4 | **85.94** | 96.82 | **79.17** | 0.00 |
|  | 1.0 | 83.13 | **98.41** | 74.17 | 8.08 |
| 36681FFPE | 0.4 | **85.75** | 98.38 | **91.00** | 4.90 |
|  | 1.0 | 81.31 | **99.05** | 85.50 | 5.22 |

Including the miRNA robustness score *DC*_Δ_ in the optimization process (*w* < 1.0) allowed to achieve higher accuracy on the validation (improvement for *BACC_cv_* ⩾ 0.46%, ⩽ 4.44%) and testing data (*BACC_te_* ⩾ 0.94%, ⩽ 7.25%) for all presented data sets. On the training data classifiers optimized using solely BACC performed better (difference in *BACC_tr_* ⩾ 0.68%, ⩽ 2.63%) for three of the data sets (22058, 36681-FF and 36681-FFPE). However, the accuracy of classifiers drops significantly when evaluated on the holdout data. This shows that training classifiers using the multi-objective *DC_score_* including the robustness score *DC*_Δ_ allows decreasing the generalization error by selecting miRNAs that display more reliable regulation patterns. Also, the standard deviation of *BACC_te_* is significantly higher for *w* = 1 (improvement for *σ_te_* ⩾ 0.32%, ⩽ 8.08%). The predictions of classifiers optimized taking the *DC*_Δ_ into account seem to increase the stability of the results.

For 22058 data set the best validation BACC was achieved for the identical weight as in the parameter tuning presented in the previous section (0.2). For 10694, 36681-FF and 36681-FFPE data sets the best performing weights were 0.9 (BACC_cv_ = 87.60%), 0.8 (BACC_cv_ = 86.16%) and 0.7 (BACC_cv_ = 87.75%), respectively. Note that that previously only approx. 10 different parameter sets were tested for each weight. Thus, the increased number of tested sets (100 per weight) may allow capturing better parameters for particular weights. Also, the *α* parameter was fixed to the values presented in previous section. Thus, the results may be further optimized.

#### 4.2.3 Comparison to SC classifiers

To design single-circuit classifiers we employed the approach described by Becker *et al*. (2018). The method allows optimizing two objectives according to the following hierarchy using the Answer Set Programming solver: (i) the numbers of errors, namely false positives and false negatives and (ii) the simplicity of the classifier. To design single-circuit classifiers we used the strategy applied for real-world case studies by Becker *et al*. (2018). For all data sets, we employ parameters that correspond to the classifier constraints proposed by Mohammadi *et al*. (2017). The method returns all globally optimal solutions that satisfy the pre-defined constraints. We recorded all the best solutions returned for each training data set and evaluated them on the corresponding testing data set using BACC. This allows directly comparing the performance of both classifier designs. In Table 7, we present average performance measures obtained for single-circuit classifiers and distributed classifiers.

**Table 7.** Performance measures for single-circuit (SC) and distributed classifiers (DC) for four cancer data sets: BACC_tr_ - average training BACC, BACC_te_ - average testing BACC, *σ*_te_ - standard deviation for BACC_te_, TPR - True Positive Rate, TNR - True Negative Rate, 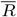 - average number of rules, 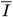 - average number of inputs.

| dataset | method | $BACC_{tr}$ | $BACC_{te}$ | $\sigma_{te}$ | TPR | TNR | $\bar{R}$ | $\bar{I}$ |
| --- | --- | --- | --- | --- | --- | --- | --- | --- |
| 22058 | DC | 97.37 | **97.50** | 0.00 | 100.00 | 95.00 | 2.0 | 3.0 |
|  | SC | **100.00** | 96.25 | 3.23 | 93.75 | 98.75 | - | 4.0 |
| 10694 | DC | **94.35** | **85.00** | 1.88 | 80.63 | 89.38 | 5.0 | 7.5 |
|  | SC | 91.13 | 84.38 | 3.61 | 87.50 | 81.25 | - | 5.0 |
| 36681FF | DC | 96.93 | **78.75** | 1.25 | 81.67 | 75.83 | 4.0 | 6.2 |
|  | SC | **98.86** | 76.67 | 3.73 | 81.67 | 71.67 | - | 5.0 |
| 36681FFPE | DC | **96.89** | **93.50** | 3.91 | 89.00 | 98.00 | 3.7 | 6.1 |
|  | SC | 93.24 | 85.56 | 3.91 | 75.56 | 95.56 | - | 5.0 |

Distributed classifiers performed better than single-circuit classifiers for two data sets on the training data (10694 and 36681-FFPE). For 22058 and 36681-FF data sets SCs achieved higher *BACC_tr_*. Evaluation on the holdout data shows that DCs performed with higher accuracy than SC classifiers (improvement in *BACC_te_* ⩾ 0.62%, ⩽ 7.94%). This demonstrates that DCs may be more robust facing novel information. Also, the *σ*_te_ is significantly lower for the distributed classifiers indicating that one may expect more reliable predictions. The increased performance may come from different features of the presented framework. First, the architecture of DCs allows for collective decisions. Second, we apply the robustness measure *DC*_Δ_ to increase the quality of miRNAs inputs. Last, we propose a framework that allows training classifiers to perform with the highest accuracy on holdout data.

Sensitivity (TPR) and specificity (TNR) measure the percentages of correctly classified cancerous and non-cancerous samples, respectively. For two of the data sets (10694 and 36681-FF) the SCs identify the cancerous samples better or equally well. For the remaining data sets (22058 and 36681-FFPE) the sensitivity of DCs is higher. This estimates the effectiveness of the classifier circuits to activate the drug production in diseased cells. However, specificity (TNR) of classifiers found by our method is increased in comparison to SC classifiers for the most of data sets, in particular for the 10694 data set. DCs capture truly negative samples more often, which may be important in terms of the application. The drug is released in each healthy cell that may be incorrectly classified as cancerous, causing its apoptosis and increasing the potential toxicity of the method.

For most of the data sets the best performing *α* equals 0.45 and the average size of the classifiers among the data sets is ⩾ 3.7, i.e., approx. half of the rules must agree to classify a cell as cancerous. For the 22058 data set *α* equals 0.25 and the average size of trained classifiers is 2.0. For classifiers consisting of only 2 rules *α* = 0.25 and *α* = 0.45 results in the same threshold, i.e., at least one of two rules must classify a cell as cancerous. This suggests that *α* of intermediate values (around 0.50) may be the most adequate for the real-world data. We analyze the structure of classifiers further in the following section using an exemplary classifier.

The DCs trained for 22058 data set achieve the highest performance among all data sets. The tuned weight *w* is lower than for other data sets. We compared Δs calculated for all miRNAs in the presented data sets using the training fraction. For 22058 data set the miRNAs achieve the highest Δs among all data sets (Δ*_avg_* = 0.64, Δ_*σ*_ = 0.10, Δ*_max_* = 0.93). The data set includes very robust candidates for classifier inputs (Δ*_max_* = 0.93). However, the differences in the robustness of miRNAs are large (Δ_*σ*_ = 0.10). Thus, the contribution of the *DC*_Δ_ score to the optimization process may be higher. In comparison, for 10694 data set the average Δ as well as the standard deviation are lower (Δ*_avg_* = 0.58, Δ_*σ*_ = 0.05, Δ*_max_* = 0.66). Also, the best potential candidate miRNA achieves substantially lower Δ*_max_*. Here, the contribution of *DC*_Δ_ is decreased (w = 0.8). These relationships should be further investigated.

In terms of simplicity single-circuit classifiers consist of a lower number of inputs on average, except for 22058 data set. However, the DCs consist of a few simple single-circuit classifiers. Note, for both designs the maximal number of inputs in the classifier was equal (10 inputs). Although the accuracy of classifiers is the most important criterion in the optimization of cell classifiers due to the application, the simplicity of the circuits influences the feasibility of their assembly. Thus, one may take it into account while designing cell classifiers.

#### 4.2.4 HCC classifier evaluation

Here, we present an exemplary distributed classifier found for the HCC data set GSE10694 that achieved the highest *DC_score_*. The classifier consists of 5 single-circuit classifiers: three 2-input (miR-224 AND miR-221, miR-224 AND NOT hsa-miR-99a, miR-93 AND NOT miR-221) and two 1-input circuits (miR-221, NOT miR-422a). The classifier was found for *α*=0.45, i.e., at least 2 rules must output 1 (cancerous) to classify the sample as cancerous.

We reviewed the literature to evaluate the viability of proposed miRNA inputs. miR-221, miR-93 and miR-224 are described as up-regulated (Liu *et al*., 2016; Ji *et al*., 2017; Hung *et al*., 2018) and miR-99ais described as down-regulated in HCC cells (Zhang *et al*., 2014). Although miR-221 is described as highly expressed in HCC tissues, the miRNA appears non-negated in 2 rules and once negated together with non-negated miR-93 (miR-93 AND NOT miR-221). Such expression pattern (miR-93 is up-regulated and miR-221 down-regulated) exists in the training data. For 13.71% of training samples the miR-93 is up-regulated and miR-221 down-regulated simultaneously (58.82% cancerous and 41.18% non-cancerous samples). For those samples, the miR-93 AND NOT miR-221 classifier outputs 1, i.e., classify them as cancerous. However, after the evaluation of the rest of rules on the same set of samples 100% non-cancerous and 90% cancerous of them were correctly classified, i.e., the DC would activate only in the cancerous cells according to the given decision threshold (miR-93 AND NOT miR-221 outputs 1 simultaneously with either NOT miR-422a or miR-224 AND NOT hsa-miR-99a, or both of them). This demonstrates the ability of DCs to capture rare or more complex expression patterns. In terms of the application, where the heterogeneity of cancer cells poses the major challenge, such ability may be a valuable feature.

## 5 Discussion

In this study, we demonstrated the potential of distributed classifiers as a new theoretical design of cell classifier circuits. We presented a comprehensive framework to design and evaluate DCs using simulated and real-world cancer data sets. We proposed a classifier-independent data simulation method that allows testing various architectures of classifiers against different types and intensities of noise. Also, we employed a discretization method that allows assessing the quality of miRNAs in terms of their robustness. Based on that, we developed a multi-objective function that optimizes two essential criteria, namely the accuracy of DCs and the robustness of employed miRNAs, to improve the performance of distributed classifiers. Further, we proposed an extensive optimization and evaluation strategy that allows to train and test classifiers against novel information. Finally, we compared the performance of distributed and single-circuit classifiers. The cancer case studies demonstrate the ability of DCs to outperform SC classifiers in terms of accuracy and robustness. Last, we showed that distributed classifiers include miRNAs that follow regulation patterns described in the literature. However, the DCs have also an ability to capture rare and complex patterns that may carry valuable information in terms of cancer cell heterogeneity.

Distributed classifiers demonstrated their high performance in several synthetic and cancer studies presented in this work. The increased performance of DCs partially derives from their architecture that allows capturing rare and more complex patterns existing in the data. Further, the proposed framework comprises an extensive optimization and evaluation scheme also significantly contributing to the classifier’s accuracy. First, the DCs have built-in robustness to noise and novel information due to the employed multi-criteria optimization function. Second, the proposed tuning and training scheme provides parameters suitable for a given data set to perform well facing novel information.

The method proposed by Becker *et al*. (2018) does not employ additional information coming from the discretization process. However, the approach has the ability to search through the entire space of solutions in a feasible time and does not require tuning of parameters. Although the method allows for finding globally optimal solutions, the single-circuit classifiers seem to over-fit the training data. Thus, the performance of SC classifiers on holdout data is decreased in comparison to DCs. The computational method proposed in Mohammadi *et al*. (2017) takes into account the circuit’s robustness by additionally optimizing so-called classification margins between the circuit output concentrations. However, the model of single-circuit classifiers, as well as the employed optimization criteria, are not directly applicable to the design of distributed classifiers. Also, in terms of binary classification, the method proposed by Becker *et al*. (2018) outperformed the previous approach.

Discretization of the expression data is an important pre-processing step that allows selecting relevant miRNA-candidates for the classifier’s inputs. To discretize each of the data sets we applied default parameters proposed by Wang *et al*. (2014). However, the parameters may be also adjusted for a given data set. Increasing values of the binarization parameters *α_bin_* and *λ_bin_* results in a more strict filtering procedure. For the default parameters, the number of miRNAs after the filtering was significantly decreased for all of the data sets. However, one may consider tuning the binarization parameters to improve the miRNA selection. In this work, we do not filter miRNAs by sequence similarities or applying other additional constraints (Mohammadi *et al*., 2017). However, such a feature may be introduced in the future development of the workflow.

We developed a multi-objective function that allows combining two optimization criteria, the accuracy of the classifier and the robustness of employed miRNAs. The importance of particular criteria may be defined by the weight. Using real-world data we demonstrated the ability of the robustness score to increase the performance of distributed classifiers. However, the weight seems to be data-dependent, e.g., the high variance between the miRNA robustness in a given data set may increase the value of the robustness score. The influence of the data-related features on the importance of both criteria should be further investigated.

In this work, we proposed an exemplary architecture of distributed cell classifiers. The constraints applied to DC’s design are based on the ones proposed for single-circuit classifiers by Mohammadi *et al*. (2017). However, the proposed architecture may be extended by employing various rule designs such as 3-input disjunctions or conjunctions or by allowing for ten 1-input rules in the DC. Such extensions may allow improving the classification performance of DCs. Thus, different architectures should be further investigated. So far, distributed cell classifiers were not evaluated in cell environments in terms of cancer classification. However, multi-circuit programs implemented in living cells were successfully studied before (Moon *et al*., 2012). Also, computing systems with increasing complexity operating in living cells seem to become available (Yehl and Lu, 2017; Lapique and Benenson, 2018). As research in synthetic biology moves in the direction of multi-device computing systems implemented in living cells presented work may provide valuable insights into their design.

## Acknowledgements

We would like to thank to Jakub M. Bartoszewicz and Bernhard Y. Renard (Hasso Plattner Institute, Germany) for their valuable comments, in particular, to JMB for constructive discussions on the testing scheme design. This work was supported by the BMBF-funded de.NBI Cloud within the German Network for Bioinformatics Infrastructure (de.NBI) (031A537B, 031A533A, 031A538A, 031A533B, 031A535A, 031A537C, 031A534A, 031A532B).

